# Predicting A/B compartments from histone modifications using deep learning

**DOI:** 10.1101/2022.04.19.488754

**Authors:** Suchen Zheng, Nitya Thakkar, Hannah L. Harris, Megan Zhang, Susanna Liu, Mark Gerstein, Erez Lieberman Aiden, M. Jordan Rowley, William Stafford Noble, Gamze Gürsoy, Ritambhara Singh

## Abstract

Genomes fold into organizational units in the 3D space that can influence critical biological functions. In particular, the organization of chromatin into A and B compartments segregates its active regions from inactive regions. Compartments, evident in Hi-C contact matrices, have been used to describe cell-type specific changes in the A/B organization. However, obtaining Hi-C data for all cell and tissue types of interest is prohibitively expensive, which has limited the widespread consideration of compartment status. We present a prediction tool called **Co**mpartment prediction using **R**ecurrent **N**eural **N**etwork (CoRNN) that models the relationship between the compartmental organization of the genome and histone modification enrichment. Our model predicts A/B compartments, in a cross-cell type setting, with an average area under the ROC curve of 90.9%. Our cell type-specific compartment predictions show high overlap with known functional elements. We investigate our predictions by systematically removing combinations of histone marks and find that H3K27ac and H3K36me3 are the most predictive marks. We then perform a detailed analysis of loci where compartment status cannot be accurately predicted from these marks. These regions represent chromatin with ambiguous compartmental status, likely due to variations in status within the population of cells. These ambiguous loci also show highly variable compartmental status between biological replicates in the same GM12878 cell type. Finally, we demonstrate the generalizability of our model by predicting compartments in independent tissue samples. Our software and trained model are publicly available at https://github.com/rsinghlab/CoRNN.

## INTRODUCTION

The physical organization of DNA inside the cell nucleus directly impacts the function and biology of the genome. DNA organization has been implicated in numerous biological processes from differentiation to oncogenesis (Zheng and Xie, 2019). Genome-wide chromosome conformation capture (Hi-C) and related techniques enable the characterization of this organization by capturing the long-range pairwise interactions among different genomic elements (Dekker et al., 2002; Lieberman-Aiden et al., 2009; Duan et al., 2010; Montefiori et al., 2016). Recent advances in the Hi-C method provide a more refined view of the relationship between genome organization and epigenomic marks (Rao et al., 2014). Detailed analyses of Hi-C interaction matrices revealed 3D structural units of chromosomes that accommodate spatial clustering of regulatory elements and identified transcription factors that are important for several cellular activities (Phillips-Cremins et al., 2013).

Hi-C data revealed that the genome is organized into two distinct compartments, labeled “A” (active) and “B” (inactive). Each of these compartments corresponds to distinct properties of the associated genomic regions. For example, there are preferential interactions within compartment types, such that loci in the A compartment tend to interact with loci in the same compartment. Compartments are also found to correlate with histone modification patterns as, for example, it was shown that there is a high concordance between ChIP-Seq signal of active histone mark enrichments such as H3K4me1 in regions that are located in A compartments (Lieberman-Aiden et al., 2009). A/B compartment boundaries are typically identified by applying principal components analysis (PCA) to the correlation matrix obtained from the Hi-C interaction frequency matrix, in which the sign of the first principal component corresponds to the A/B compartments. Conventionally, loci associated with A compartments (active) are assigned positive values, while those in B compartments (inactive) are assigned negative values in the eigenvector.

While Hi-C is a powerful experimental technique to detect chromosomal compartments, the high cost and technical difficulties make obtaining Hi-C data for many different cell lines and types challenging. Therefore, predicting these organizational units of chromosomes via more abundant data types, such as ChIP-Seq, can remedy the lack of Hi-C data. Furthermore, such prediction methods can provide insight into the interplay between the 3D organization of the chromosome and its 1D activity level. Therefore, it is crucial to 1) find ways to infer cell-type specific compartments without needing Hi-C data generation and 2) discover relationships between compartments and chromatin marks to better understand the connections between the spatial organization and biology of the genome.

Previous methods have used epigenetic signals like DNA methylation (Fortin and Hansen, 2015; Jenkinson et al., 2017; Raineri et al., 2018; Al Bkhetan and Plewczynski, 2018) to capture such relationships. For example, Fortin and Hansen (2015) used the eigenvectors calculated from correlation matrices of DNA methylation experiments and reported correlation values of ~ 0.56 – 0.71 with the A/B compartments. Jenkinson et al. (2017) showed that entropy blocks calculated from DNA methylation data correspond well to the topologically associating domain (TAD) boundaries obtained from Hi-C data. Raineri et al. (2018) used a linear regression model to predict compartments from GC-content and DNA methylation experiments and reported a mean absolute error of 0.9. Al Bkhetan and Plewczynski (2018) used a random forest model to predict contact loops, obtained from ChIA-PET experiments, transcription factors, and histone modification experiments. They reported accuracy of 0.87 for their model (3DEpiLoop). However, none of these existing methods explored the specific relationships between the histone modifications signals and A/B compartments of the genome.

We hypothesize that since A/B compartment assignments are proxies for genome activity, similar information can be inferred from analyzing the histone modification data obtained from ChIP-Seq experiments. To this end, we propose a deep learning framework for **Co**mpartment prediction using **R**ecurrent **N**eural **N**etworks (CoRNN) to predict chromosome compartments using histone modification ChIP-Seq data (Fig. 1). Recent studies have demonstrated the effectiveness of neural networks in sub-compartment calling using Hi-C data (Ashoor et al., 2020). Here we apply neural networks for predicting the compartments without Hi-C data.

**Figure 1.**
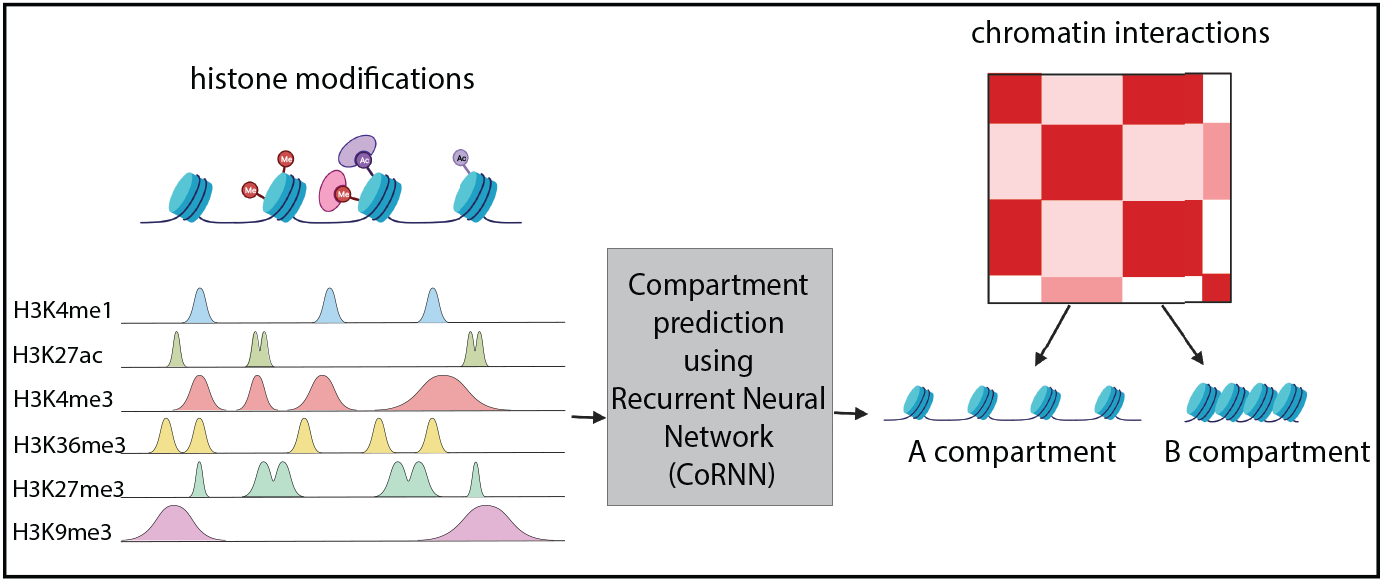
Overview: The A/B compartment prediction task is formulated as a binary classification problem. We use six histone modification ChIP-Seq experiments as our inputs: H3K4me1, H3K27ac, H3K4me3, H3K36me3, H3K27me3, and H3K9me3. Our framework, Compartment prediction using Recurrent Neural Networks (CoRNN), uses a recurrent neural network to model the input features and mean compartment values for cell lines in the training set and predict A/B compartments for the cell lines in the test set.

We carefully selected three baselines to benchmark the performance of CoRNN. It was previously shown that in genomic prediction tasks, a machine learning model might falsely appear to perform well by effectively memorizing the average activity associated with each locus across the training cell types (Schreiber et al., 2020b). Therefore, first, we use the mean compartment value baseline (or the “mean baseline”) that predicts the compartment assignment of a genomic region based on the average compartment values across cell lines in the training set. Since most compartments are conserved across cell types (Lieberman-Aiden et al., 2009), this baseline is quite difficult to beat. The second baseline is a random forest model (similar to Al Bkhetan and Plewczynski (2018)) that uses the mean and standard deviation of six histone modifications and the mean compartment value as model input. Finally, the third baseline is a logistic regression model (similar to Raineri et al. (2018)), which uses the same input as the random forest model. CoRNN predicts the compartment assignments better than these three competing baselines, yielding an improvement of 10% in AUROC score on average for all cell lines.

Next, we looked at highly variable regions that CoRNN predicted correctly and the mean baseline predicted incorrectly, and vice versa. We show that over 90% of these “disagreement” regions overlap with known candidate cis-regulatory elements. This overlap is higher than the overlap found for the regions that are correctly predicted by the mean baseline but missed by CoRNN (~ 30%). This result indicates that the highly variable regions mispredicted by the mean baseline are biologically important and are correctly labeled by CoRNN. We also performed a perturbation analysis to identify the histone modifications that are highly predictive of A/B compartment status. We found that H3K27ac and H3K36me3 are the most informative histone marks for CoRNN. Furthermore, we investigated the genomic regions for which CoRNN predictions and the Hi-C eigenvector values did not match. We find that these difficult-to-predict regions correspond to highly ambiguous compartment scores that vary between different Hi-C biological replicates of the same cell line, GM12878 (Rao et al., 2014). Finally, we tested CoRNN with two independent test datasets from the human muscle and colon tissues to assess the model’s generalizability. We show that CoRNN outperforms the mean baseline by 13.9% for these previously unseen datasets.

Overall, this study and our new framework CoRNN enable assigning A/B compartments to genomic regions for cell lines and tissues with no available Hi-C data. Our perturbation analysis shows that highly accurate predictions can be made even when using only a couple of histone modification ChIP-Seq data sets, thereby enabling inference of large-scale genome organization from experimental data that is easier and cheaper to obtain than Hi-C.

## METHOD

### Data Preprocessing

For training the model, we selected Hi-C and histone modification ChIP-seq experiments for six cell lines: NHEK (normal human epidermal keratinocytes), IMR90 (normal human lung fibroblasts), HMEC (human mammary epithelial cells), GM12878 (human lymphoblastoid cells), K562 (myelogenous leukemia cells), and HUVEC (human umbilical vein endothelial cells). We predict the A/B compartments for each cell line (test set) by training the CoRNN model on the other five cell lines (training set). The model takes two inputs: histone modification signals of the test cell line and mean compartment values of the training cell lines. To generate the input using histone modification signals, we divided the chromosome into 100 kbp regions for each cell line and binned each region into 100 bins of size 1000 bp. For each bin, we calculated the average histone modification ChIP-seq signal. We chose the following six histone modification marks: H3K4me3, H3K4me1, H3K27ac, H3K36me3, H3K9me3, and H3K27me3. These marks were selected because they are consistently available across most of the six cell lines.

Next, we obtained the A/B compartment values by calculating the first-order eigenvectors of the Hi-C matrix (at 100 kbp resolution) for each cell line. We formulate the A/B compartment prediction as a binary classification problem. Therefore, we assigned output labels 1 (A compartment) and 0 (B compartment) to positive and negative compartment values, respectively.

We also eliminated regions with a missing compartment value and imputed input for regions with missing histone modification signals. For example, for NHEK, the H3K27me3 and H3K36me3 experiments were missing. Similarly, for GM12878, the H3K9me3 experiment was missing. We imputed the missing histone modification values using the average ChIP-Seq signal across other cell lines.

Finally, we included the histone modification marks and compartment values from the Colon and Muscle tissue samples taken from a consented individual (accession code ENCDO845WKR) in the ENCODE portal (http://entex.encodeproject.org/) as an independent test set to demonstrate the generalizability of our model.

### Input and output formulation for the prediction task

Fig. 2 shows an example input sample (representing a 100 kbp genomic region) denoted as a matrix 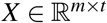. Here, *m* = 6 denotes the number of histone marks, and *t* = 100 are the number of genomic bins. We input matrix *X* and scalar *c*, which is the mean compartment value for the training cell lines, for each genomic region and predict its compartment. The output *y* ∈ [0,1] represents the binarized compartment value for the input genomic region.

**Figure 2.**
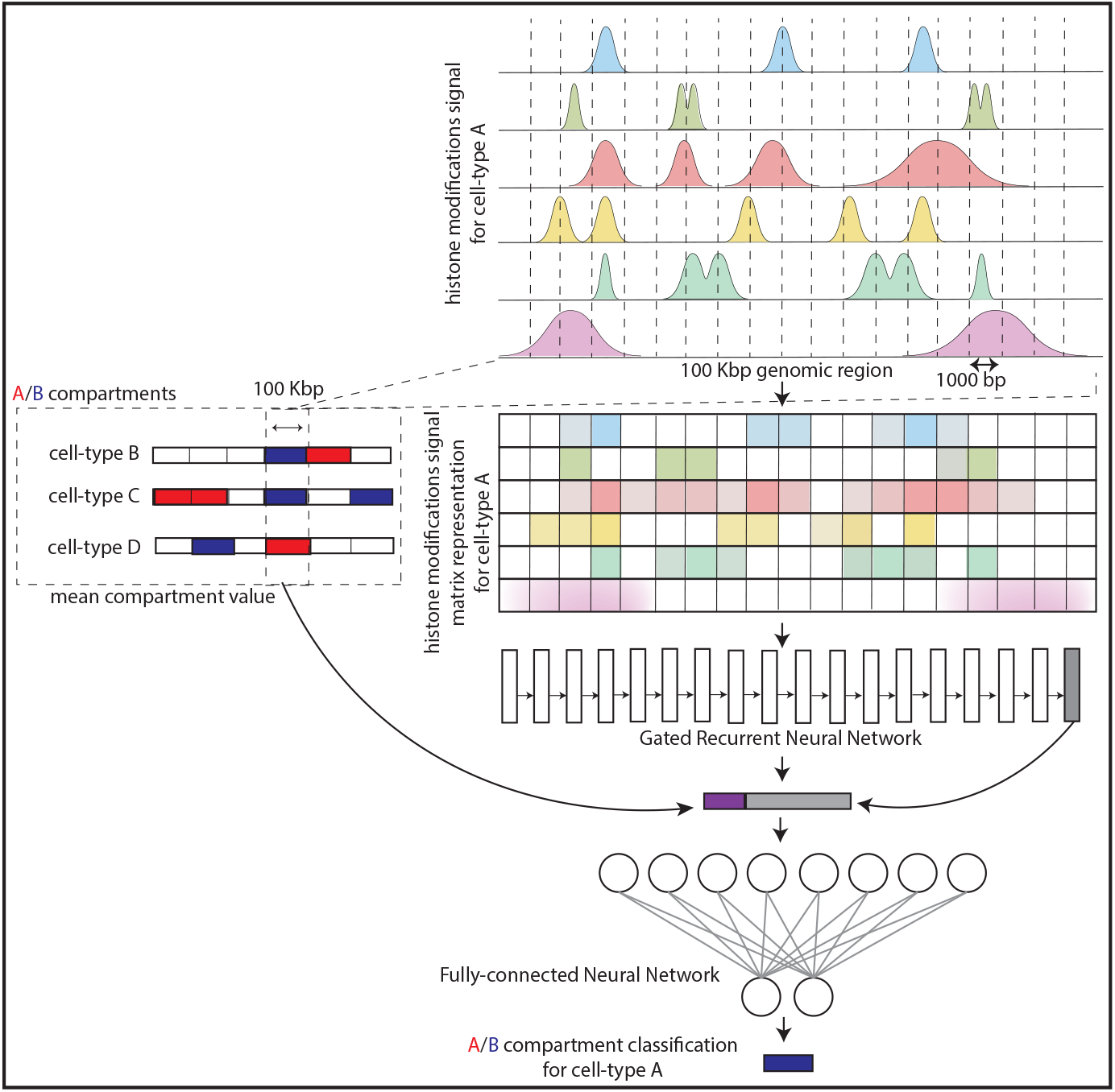
CoRNN architecture: The main input into the model is a matrix 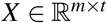. Here, *m* = 6 denotes the number of histone marks, and *t* = 100 are the number of genomic bins representing a 100kbp genomic region. The model consists of gated recurrent unit (GRU) layers to capture the sequential information of the histone modification signals across the genomic region. The output of the GRU is then concatenated with the mean compartment value *c* for the training cell lines and fed into fully connected layers. The output *y* ∈ [0,1] represents the binarized compartment value for the input genomic region.

### CoRNN architecture

CoRNN is an end-to-end A/B compartment prediction model (Fig. 2). It consists of three main components:

#### Gated recurrent units (GRUs)

Gated recurrent units (GRUs) are a variation of the traditional recurrent neural network (Cho et al., 2014). GRUs can capture long-range sequential information from the input samples. We also tested a convolutional neural network (CNN) as an architecture choice, but it did not perform as well as the GRU (Supplementary Fig. 8). Therefore, in our setting, we hypothesize that a GRU layer effectively models the sequential dependency of the histone marks in consecutive bins across the genome resulting in better performance.

Given our input matrix *X*, GRUs take in one input column *x_t_* (with all six histone marks) at a time. Together with the hidden state *h*_*t*–1_ from the previous time step, GRUs generate the current hidden state *h_t_* as the input to the next time step. More specifically, GRUs first calculate the update gate *z_t_* for time step *t* using

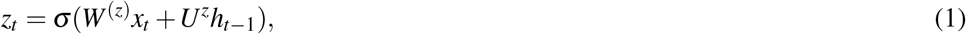

where current input *x_t_* is multiplied by its weight *W*^(*z*)^ and hidden state *h*_*t*–1_ from the previous time step is multiplied by its weight *U^z^*. These two values are added and inputted to a sigmoid activation function (Equation 2) to constrain the result between 0 and 1. The update gate function acts as the long-term memory of the network. It determines how much past information will need to be passed down to the next step:

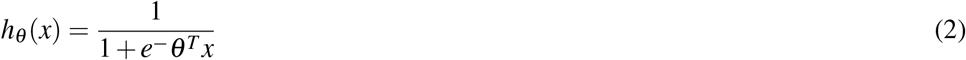

GRUs also have a reset gate to determine the short-term memory of the network, that is, how much information to discard, using the following formula:

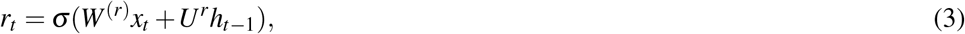

Next, GRUs determine the current memory content by applying the output of the reset gate *r_t_* to the hidden state from the previous time step *h*_*t*–1_ (Equation 4). This step uses an element-wise product between *r_t_* and *Uh*_*t*–1_. The current input *x_t_* is multiplied by weight *W*. These values are added together and inputted to the *tanh* activation function:

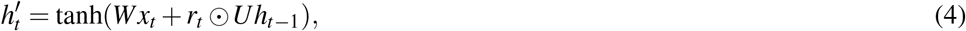

Finally, GRUs calculate the *h_t_* using the following formula:

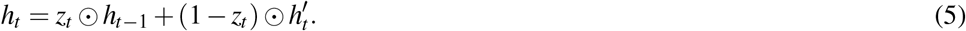

When the GRU sets *z_t_* close to 1, it will retain most of the information from the previous hidden state *h*_*t*–1_. Since (1 – *z_t_*) will be close to 0, the model will ignore most of the current content from 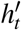.

We use GRUs to learn the representation of the histone modification signals. The number of GRU layers and the size of the hidden units are hyperparameters of the model (Supplementary Table 1). When incorporating multiple layers, only the first GRU layer takes the original histone modification signals as input. The subsequent layers take the hidden state outputs from the previous layer as input. The output of the last hidden unit, *h*_100_, of the final GRU layer, concatenated with mean compartment value *c*, goes into the next component of the model, the fully connected network.

#### Fully connected network (FCN)

This network consists of two fully connected layers. It takes the last hidden state of the GRU and the mean compartment value as inputs and generates an output vector of size two. By concatenating the mean compartment value to the GRU’s output, we enable CoRNN to leverage information from histone modification signals and the compartment consensus of other cell lines in the training set. This operation results in a vector of size *h*_100_ + 1 as the input to the fully connected network. Here, || represents concatenation, c represents the mean compartment value. *W*_1_ and *b*_1_ represent the learnable weight and bias parameters of the first fully connected layer, and *W*_2_ and *b*_2_ represent the learnable weight and bias parameters of the second fully connected layer. Therefore, the output of this network can be written as

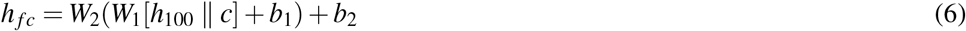

#### Softmax function

Finally, a softmax function is applied to *h_fc_*. We formulate the A/B compartment prediction as a binary classification task with classes *y* ∈ {0,1}, corresponding to whether a chromosome region is in A (active) compartment (*y* = 1) or in B (inactive) compartment (*y* = 0). The softmax function takes in the output value from the fully connected network and computes the probability of each class y. We use the cross-entropy loss for the predicted probability of the true label to train the model’s weights.

### End-to-end training

Out of the six selected cell lines, we iteratively choose one cell line as the test set and the other five cell lines as the training set. The mean compartment value is calculated by averaging the compartment values across all training cells on the same chromosome and region. We present our cross-validation scheme in Supplementary Fig. 9. For example, if IMR90 is the test cell line, we use GM12878, K562, NHEK, HMEC, and HUVEC as the training set and perform hyperparameter selection using five-fold cross-validation. For each cross-validation fold, we select one cell line from the training set as validation for the current fold. Then we train the model on each fold and obtain the average validation performance from the five folds. Finally, the best average validation performance model is used to make the test cell line predictions. When training CoRNN, we hold out the test cell (e.g., IMR90) from all aspects of the process and use it solely to report the performance of the final model. While training CoRNN, we performed hyperparameter tuning over the following grid of values to pick the best model architecture: the size of hidden state ∈ {32,64,128} and the number of GRU layers ∈ {1,2,3,4}.

## EXPERIMENTAL SETUP

### Baseline methods

#### Mean compartment value baseline

The mean compartment value baseline (also referred to as the mean baseline) uses the average compartment values across cell lines in a training set as a proxy for the A/B compartment prediction in the test cell line. First, we binarize the compartment values to 1 or 0 based on positive and negative values. Next, we take the average of the five binarized compartment values. Since the training set comprises five cell lines, the predictions made by the mean baseline will have the following values: 0, 0.2, 0.4, 0.6, 0.8, and 1.0. A mean compartment value close to 0 or 1 for a genomic bin indicates that the compartment value is more consistent in this region across all five training cell lines, showing that this is a more conserved region. Similarly, a mean compartment value of around 0.5 means the compartment value varies across different cell lines and represents a less conserved region. Since most of the compartments are conserved across different cell lines, the mean baseline’s predictions can achieve a performance that is difficult to beat.

#### Random Forest

Al Bkhetan and Plewczynski (2018) use a Random Forest model to predict physical interaction in chromatin using a variety of histone modifications (H2AFZ, H3K27ac, H3K27me3, H3K36me3, H3K4me1, H3K4me2, H3K4me3, H3K79me2, H3K9ac, H3K9me1, H3K9me3, and H4K20me1) and transcription factors (CTCF, RNAP II, RAD12, ZNF143, SMC, and SA1). Given that we are predicting the A/B compartments using epigenomics features, we included a similar Random Forest model as one of the baselines. Its hyperparameter tuning was performed on the number of trees in the forest, the maximum depth of the tree, the minimum number of samples required to split an internal node, and the minimum number of samples required to be at a leaf node. For the model input, we calculated the mean and standard deviation for each of the six histone modification signals in the 100 kbp region for input features. Mean and standard deviation values from six histone modification signals made up an input vector of length 12. To keep this model consistent with our framework and for a fair comparison, we concatenate the mean compartment value at the end of the input vector. We also tried using all of the 6 × 100 features as input to train the Random Forest model. However, the performance of this model was worse than using mean and standard deviation values (Supplementary Fig. 10).

#### Logistic regression

Raineri et al. (2018) proposed a logistic regression framework to predict A/B compartments from GC content of the sequence and DNA methylation. Following their setup, we trained a logistic regression model to include as one of our baselines. We used the same data pre-processing as the Random Forest baseline. The input to the model was the mean and standard deviation of six histone modification signals in the 100 kbp region combined with the mean compartment value of the region across the training cell lines. We performed hyperparameter tuning of the model for the norm of the penalty (*l*1, *l*2), the *C* value (inverse of regularization strength), type of solver (*newton – cg; lb f gs; liblinear;sag, saga*), and the maximum number of iterations taken for the solvers to converge. We also tried using all of the 6 × 100 features as input to train the logistic regression model. However, similar to the Random Forest model, the performance using all 600 features was not as good as using the mean and standard deviation of the signals (Supplementary Fig. 10).

### Evaluation metrics

We trained all the models on the five cell lines and selected the best performing hyperparameters using a five-fold cross-validation scheme. We then tested the selected model on the sixth cell line. Since we formulate the compartment prediction problem as a binary classification task, we use the area under the receiver operating characteristic (AUROC) score as our evaluation metric. The AUROC score evaluates the classifier’s ability to distinguish two classes. It measures the probability that a random positive sample will be ranked higher than a randomly selected negative sample. The AUROC score ranges between 0 and 1, where values closer to 1 indicate a more successful classifier. Since the number of samples in our two classes—A and B compartments—are roughly balanced (Supplementary Table 2), our choice of AUROC score is reasonable. However, we have also included the area under the precision-recall curve (AUPRC) scores to evaluate the classification performance in the Supplementary for completeness.

## RESULTS

### CoRNN gives state-of-the-art compartment prediction performance

Fig. 3 presents the A/B compartment classification performance of CoRNN across six selected cell lines using the AUROC score. In functional genomics, especially in measurements of 3D genome configuration, most of the signal can be highly conserved across cell types (Dixon et al., 2012). For example, if we look at the correlation of compartment values among the six cell lines, we find generally high correlation values (minimum correlation is 73% and maximum correlation is 96%; Supplementary Fig. 11). This observation means that 100 kbp compartment labels across the genome are largely consistent among different cell types. Therefore, it is important to compare the predictions against the average behavior of the cell types to ensure that the model predicts cell type-specific signals (Schreiber et al., 2020b). To this end, we compared the performance of our model against the performance of the mean baseline. Our predictions are more accurate than the mean compartment values for five of the six cell lines. This result is consistent if we use the area under the precision-recall curve (AUPRC) scores as our evaluation metric (Supplementary Fig. 12). Moreover, we found that none of the other baselines could predict the labels better than the labels produced by the mean compartment values. We also include the CoRNN model performance that does not leverage the compartment values of the training data (labeled as “GRU”). The GRU model outperforms the mean baseline for only the GM12878, K562, and IMR90 cell lines.

**Figure 3.**
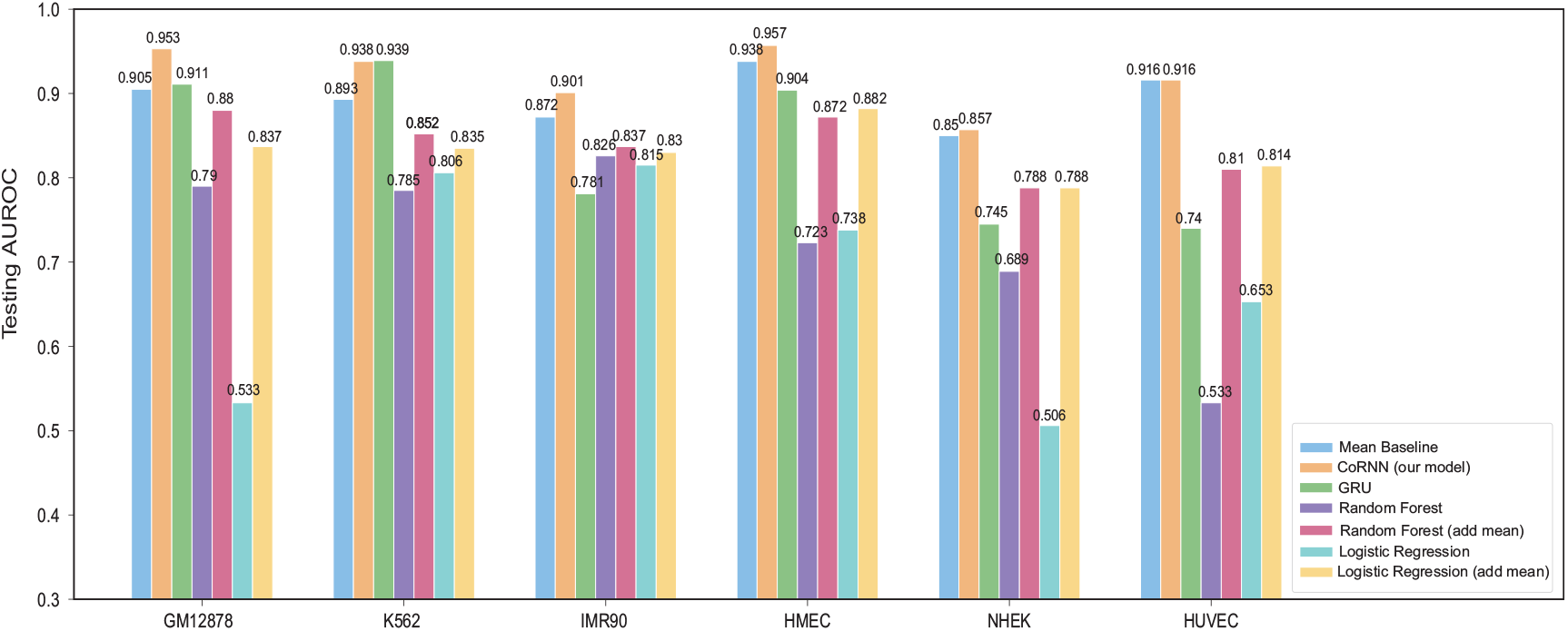
Testing results of CoRNN and baselines. Our model gives the best prediction performance across all cell lines and outperforms the mean baseline for five out of six cell lines.

### CoRNN is more accurate for regions with variable compartment values across cell lines

Many genomic regions have the same Hi-C derived compartment labels across different cell lines, while some are more variable (or dynamic) due to the cell type specificity. One way to gauge the model performance is to look at the accuracy of the predictions by categorizing the regions by their cell type specificity. For this, we divide all the genomic regions with associated compartment values into sub-groups based on their label concordance across the five training cell lines. Fig. 4 (B) plots these sub-groups on the x-axis and reports the accuracy of the mean baseline and CoRNN for these regions in the sixth test cell line on the y-axis. We cannot obtain a ranking for the mean baseline to calculate the AUROC score for this analysis as it predicts only one value (0, 0.2, 0.4, 0.6, 0.8, or 1.0) for each sub-group. Therefore, we use the accuracy metric instead. These accuracy scores have been averaged across all the test cases. A value of 0 on the x-axis represents genomic regions with B compartment labels consistent across all five cell lines, and a value of 5 represents the same for the A compartment. Similarly, 1 and 4 represent regions with A or B label concordance in four out of five cell lines and 2 and 3 for three out of five. We call the genomic regions with low A or B compartment concordance in labels across the cell lines “dynamic regions” (values 1-4). As expected, our CoRNN model exhibits a performance gain over the mean baseline for these regions, which is especially significant for regions with A compartment variability across 3 and 4 cell lines. We observe this trend because the mean baseline depends on concordance among labels to make predictions. On the other hand, our model can learn from the histone modification profiles to make more accurate predictions.

**Figure 4.**
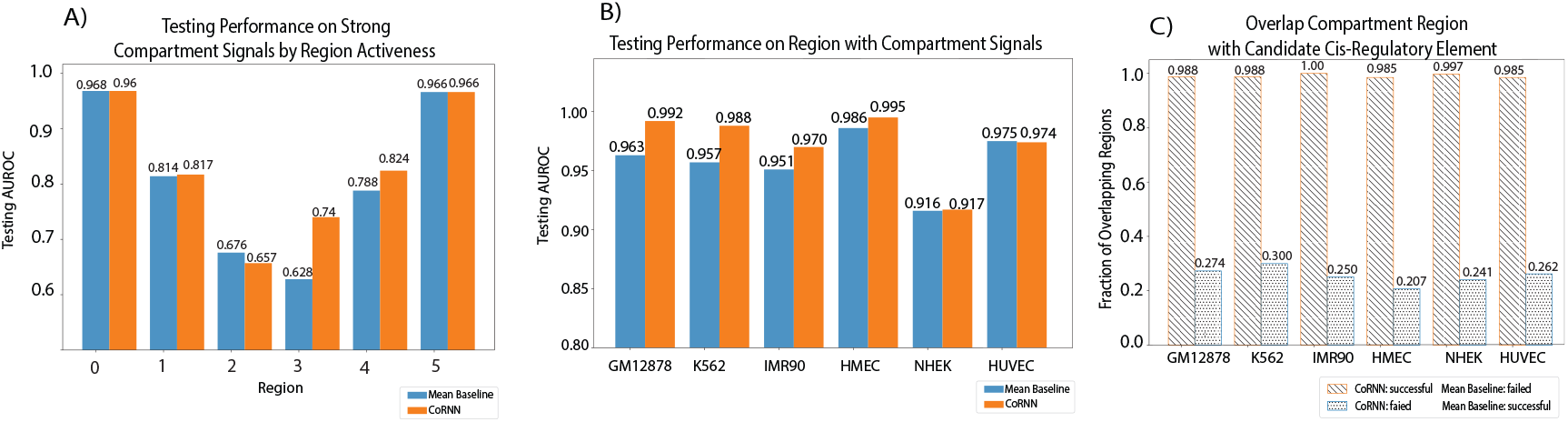
(A) Comparison of AUROC scores between CoRNN and mean baseline for all cell lines for genomic regions with strong compartment values. CoRNN predicts these regions with higher accuracy than the mean baseline. (B) Comparison of AUROC scores between CoRNN and mean baseline when the regions are categorized by consistency across cell line. CoRNN model has a performance gain over the mean baseline for these regions, especially significant for regions with A compartment variability across 3 and 4 cell lines. (C) Fraction of overlap between candidate cis-regulatory elements (cCREs) and predicted compartment for regions that are correctly predicted by CoRNN but missed by mean baseline and regions that are correctly predicted by mean baseline but missed by CoRNN. Most regions that are predicted by CoRNN but missed by mean baseline overlap with the cCREs.

### CoRNN is highly accurate for regions with high compartment values

When we look at the Hi-C data, there are regions in the genome that show weak signals (small absolute values in the eigenvector) regarding which compartment they belong to. However, it is unclear which compartments these ambiguous regions belong to and whether this is an issue with the resolution of the Hi-C data. Because we frame the A/B compartment prediction task as binary classification, we hypothesize that CoRNN would show better prediction performance for “strong” compartments (regions with high compartment values). Therefore, we select a subset of these strong compartments as those with absolute values > (*mean – std.deviation*) (Supplementary Fig. 13). In this setting, we observe a marked increase in AUROC scores for both CoRNN and the mean baseline for these strong compartments (Fig. 4(A)). In particular, our model is highly accurate at predicting these strong compartments, achieving AUROC scores *~* 0.98 for four out of six cell lines. This result suggests that CoRNN can reliably be used to predict strong compartments in a cell line using histone modification data. This property is beneficial for detecting accurate A/B compartments to analyze the 3D genome organization in a cell line without its Hi-C data.

### CoRNN correctly predicts regions with cis-Regulatory Elements

We investigated the regions that are correctly predicted by CoRNN but are missed by the mean baseline. These regions tend to be cell-type specific; hence, they cannot be assigned a compartment label based on the mean compartment values of all cell lines. We calculated the enrichment of candidate cis-regulatory element (Moore et al., 2020) (cCREs, ENCODE accession code ENCFF788SJC) on regions that are predicted by CoRNN but missed by the mean baseline and, conversely, on regions that are predicted accurately by the mean baseline but missed by CoRNN. This overlap is calculated by annotating every 100kb bin in the genome as containing cCREs if there is at least one cCRE overlapping with the bin since cCREs and compartments are in different resolution. We found that almost all of the regions (99%) that are predicted correctly by CoRNN but missed by the mean baseline overlap with cCREs. In contrast, only 20 – 30% of the complementary regions overlap with cCREs (Fig. 4 (C)). This observation holds for all cell lines, even though the predicted regions differ across cell lines. This observation suggests that most of the regions that CoRNN exclusively predicts are functionally important.

### Perturbation analysis reveals H3K27ac and H3K36me3 as the most predictive histone marks for CoRNN

Histone modifications can provide redundant information because many types of histone modifications are highly correlated with one another. We perform a perturbation analysis to determine which histone modifications have the best predictive power and are the most relevant for our accurate A/B compartment classification. We take our trained CoRNN model for each cell line and mask out all possible combinations of histone modification signals one by one by replacing the input matrix rows with zeros and recording the AUROC scores. Fig. 5 presents the performance results for the combinations of histone marks that result in a similar or higher AUROC score compared to the mean baseline. The list of histone modifications is ranked based on their frequency of occurrence for such combinations. We observe that H3K27ac and H3K36me3 are the most important histone marks for CoRNN to make accurate A/B compartment classification. H3K4me1 and H3K9me3, respectively, follow these. H3K4me3 and H3K27me3 seem to be the least relevant for the CoRNN predictions. These results align with recent studies connecting histone modifications to A/B compartments. A recent study (Sefer, 2021) found that H3K4me1, H3K9me3, and H3K27ac are some of the histone marks that are significantly predictive of most Hi–C interactions in human ES cells. Another study (Gu et al., 2021), using the ultra-resolution Hi-C contact map in lymphoblastoid cell lines (LCLs), observed the correlation of low H3K4me1 and H3K36me3 signals with the presence of discordant compartmentalization. Therefore, by modeling the relationship of histone modifications with the A/B compartments, CoRNN can capture the relevance of these marks in highlighting the properties of genomic compartmentalization. This perturbation analysis also helps us identify the minimum amount of information required for predicting A/B compartments. For example, if only H3K27ac and H3K36me3 ChIP-seq experiments are available for a given cell line, we can still make accurate A/B compartment classifications using CoRNN.

**Figure 5.**
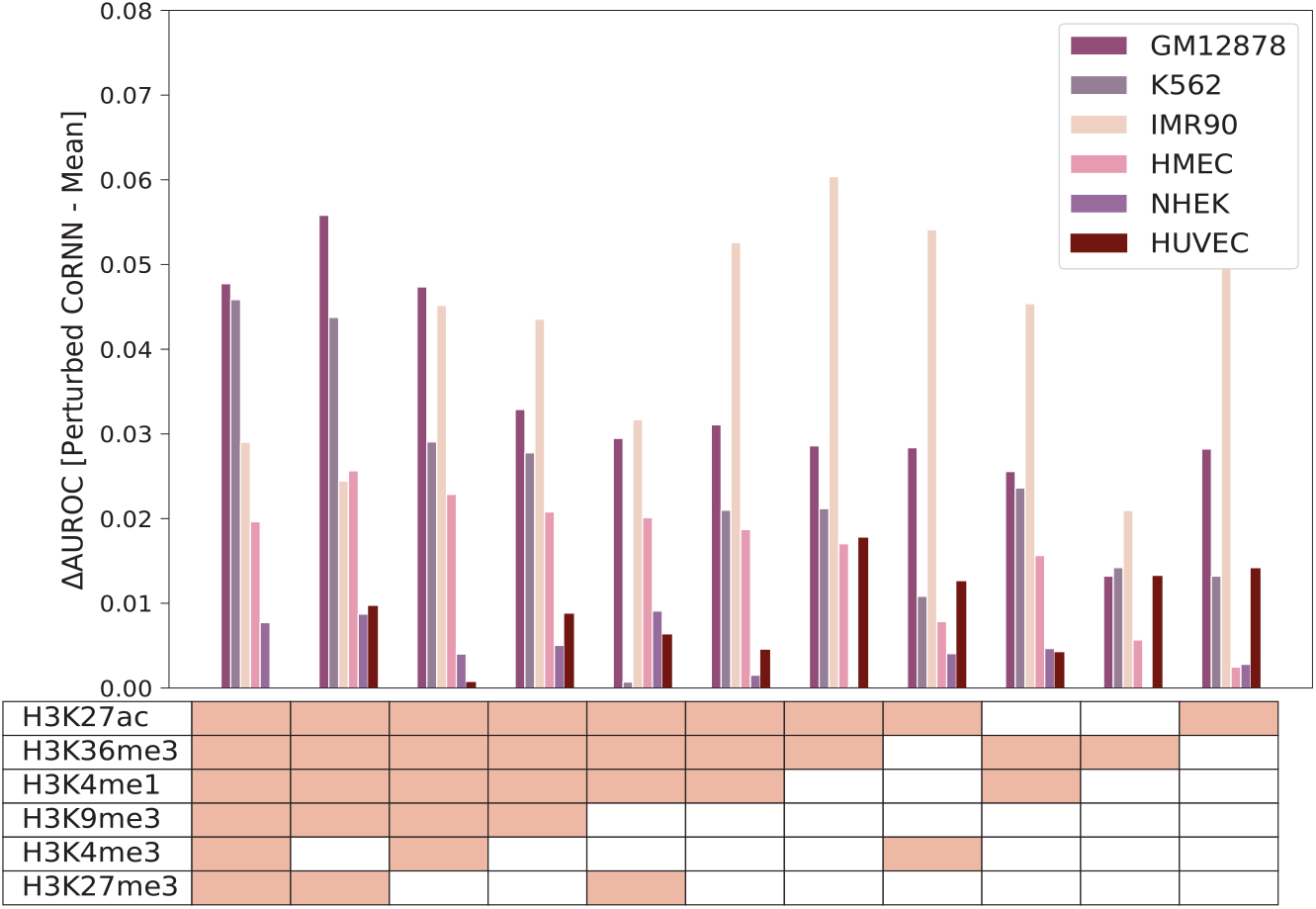
Performance results for perturbing different combinations of histone marks that result in a similar or higher AUROC score compared to the mean baseline. The list of histone modifications is ranked based on their frequency of occurrence for such combinations. We observe that H3K27ac and H3K36me3 are the most relevant histone marks for CoRNN to make accurate A/B compartment classification.

### CoRNN mispredicts regions with ambiguous chromatin marks or compartment values

We next investigate the regions that CoRNN failed to accurately predict to understand the functionality of these regions and why histone modification information is insufficient to classify their compartment values accurately. We also included Pol2Ser2 and Pol2RA signals in this analysis to understand these regions’ activity profiles better. Unsurprisingly, when we looked at the mispredictions, we found that regions predicted as A by CoRNN but B by Hi-C generally had histone marks more similar to that expected by the regions that are in the B compartment (Fig. 6(A)). However, there was an exception, histone mark H3K27me3, whose values were more similar to that of regions correctly assigned as the A compartment. This observation indicates that regions residing in the B compartment might be challenging to classify if their H3K27me3 status is similar to that of regions in the A compartment. This also means that it is difficult to predict the compartmental status of loci with both active and repressive chromatin marks, such as bivalent enhancers. Indeed, it is likely that these types of regulatory elements form unique chromatin interaction patterns (Gu et al., 2021).

**Figure 6.**
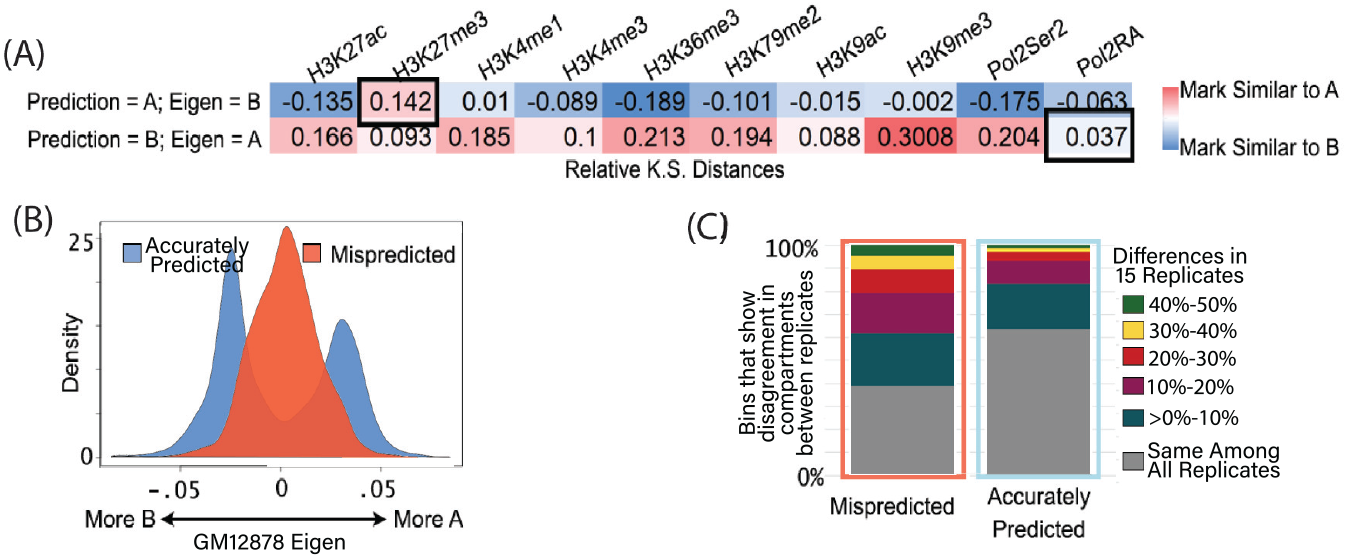
(A) Similarity of chromatin marks in mispredicted bins to that of correctly predicted A or B compartments. Values represent the subtraction of Kolmogorov-Smirnov distances for each. Black rectangles highlight marks that may have contributed to the misprediction. (B) The distribution of the eigenvector in the GM12878 map for bins that were mispredicted vs. accurately predicted. Mispredicted bins tend to have scores closer to 0, indicative of more ambiguous compartment status. (C) Examination of the eigenvector for 15 biological replicates of the GM12878 Hi-C map and the percentage of bins that show disagreement between individual maps.

In contrast, regions that were predicted as B by CoRNN but A by Hi-C had somewhat intermediate levels of active marks (Fig. 6(A)). Pol2RA levels were especially low compared to A compartment regions. Altogether, these results indicate that regions that are mispredicted by CoRNN often exhibit unusual chromatin activity mark enrichments for the compartment status designated by Hi-C. This could also be indicative of the limitations imposed by a two-state compartment model (Nichols and Corces, 2021), suggesting that sub-compartment calling can provide valuable additional information for these difficult-to-predict regions.

We further analyzed the difficult-to-predict genomic regions by examining the eigenvector values from the GM12878 Hi-C map (Rao et al., 2014). Regions that CoRNN has trouble predicting have values closer to 0 in the eigenvector, indicating a more ambiguous compartment status compared to those that are accurately predicted (Fig. 6(B)). Because the GM12878 Hi-C map represents a combination of 15 independent replicates, the ambiguous compartment status in the combined map may be due to variability between individual replicates. Using the Hi-C maps of the individual replicates, we annotated compartments from the eigenvector and examined the compartment status of each for mispredicted versus accurately predicted bins. From this analysis, we found that bins mispredicted by CoRNN often represent sites with poor agreement among replicates (Fig. 6 (C)). These analyses suggest that CoRNN is highly accurate for most regions. However, some sites are difficult to predict due to an ambiguous compartment status from unexpected chromatin marks or variability in the sampled population.

### CoRNN accurately predicts A/B compartments in independent tissue samples

Finally, we demonstrate the generalizability of a trained CoRNN model by testing it on new tissue samples. Note that the CoRNN model is trained on datasets from six cell lines, and we test it on colon and muscle tissue samples taken from a consented individual (accession code ENCDO845WKR) in the ENCODE portal (http://entex.encodeproject.org/). These datasets are entirely unseen by the model during training. We selected them because they had all the ChIP-seq and Hi-C experiments available to test and evaluate the model predictions. We present the prediction performance of CoRNN and compare it to the mean baseline in Figure 7. We observe that CoRNN predicts the A/B compartments more accurately than the mean baseline with AUROC scores of 0.72 and 0.77 and AUPRC scores of 0.79 and 0.82 (Supplementary Figure 14) for muscle and colon tissue, respectively. However, the scores, in general, are lower for the tissue samples as compared to the cell lines. We hypothesize that this observation is due to the heterogeneity of the tissue samples. Nevertheless, our results indicate that CoRNN is a useful predictive tool for new cell lines or tissues with missing Hi-C data and a better alternative than using mean values as proxies for compartment values.

**Figure 7.**
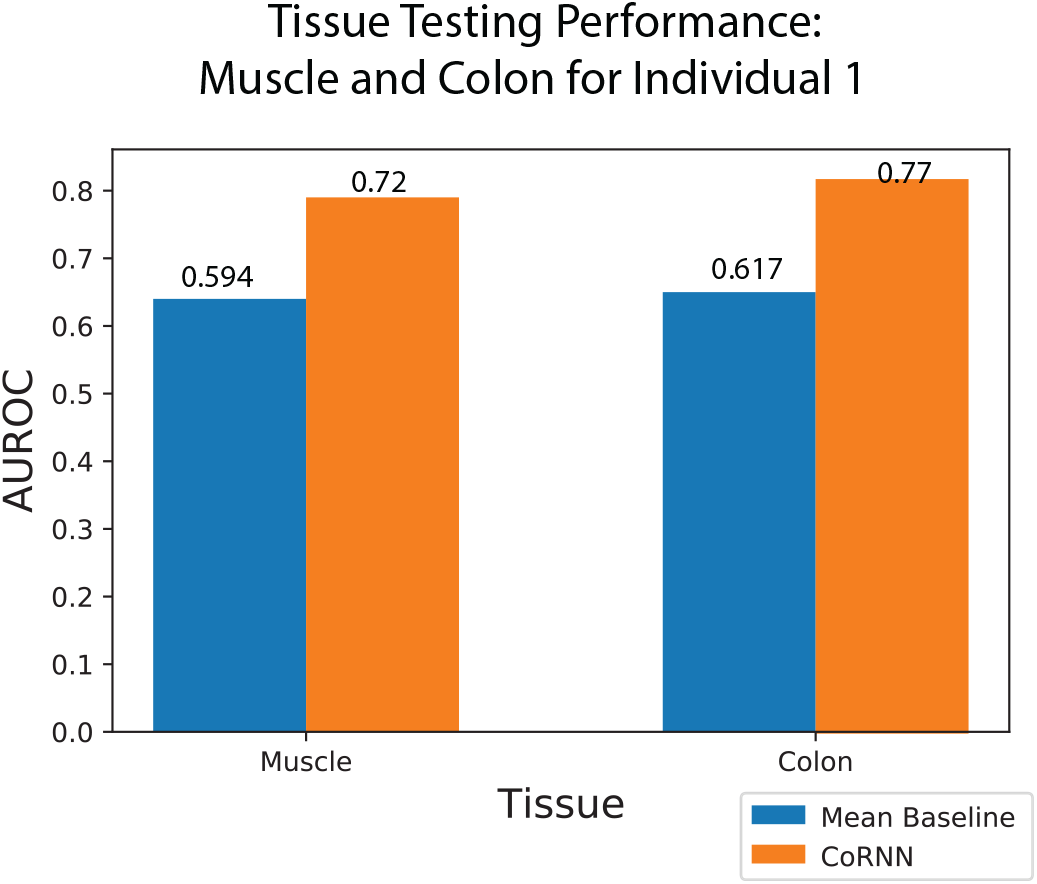
Testing CoRNN and mean baseline on human muscle and colon tissue samples. CoRNN predicts A/B compartments for both tissues with higher AUROC scores compare to mean baseline.

## DISCUSSION

We describe the CoRNN method that can take one-dimensional ChIP-seq signals from histone modification enrichment and accurately predict the chromatin compartments that otherwise require Hi-C data. This method will enable obtaining compartment designations for cell types that do not have Hi-C data available and will also allow interrogation of the relationship between the epigenomic landscape and its three-dimensional shape in the nucleus.

One of the most important benchmarks for cross-cell type predictions is to see how the predictions compare against a simple average baseline (Schreiber et al., 2020b). For example, if we were to average all the compartment scores across six cell lines, how would this average predict the compartments for any cell line? In this study, we compared all of our prediction performances against this average baseline to make sure that we predicted cell-type specific compartments.

In order to understand the histone marks that are most relevant for our predictions, we performed a detailed perturbation analysis, in which we tested all possible combinations of histone marks as features. We found that H3K27ac and H3K36me3 are the most relevant marks. H3K27ac is highly associated with transcriptional activation and is used to identify active enhancers. Therefore, we expect to see high enrichment of H3K27ac on the active non-coding genome that would be located in the active A compartments. On the other hand, H3K36me3 marks gene bodies and hence is enriched on the coding genome. Altogether, both marks have the potential to represent the entire genome and therefore are likely useful in distinguishing A/B compartments.

When we analyzed the regions that CoRNN mispredict, we found that they represent regions with unusual marks for their Hi-C annotated compartment. For example, a region that is enriched by active histone marks but labeled as inactive B compartment would be denoted as a misprediction. However, we postulate that this might also depend on the resolution of the Hi-C data. Our compartment calls are made at 100 kb resolution, which means any compartmental region that is smaller than 100 kb might be erroneously labeled with its neighboring compartment (Gu et al., 2021). Additionally, these regions have eigenvector values close to 0, indicative of some ambiguity in compartment status measured by Hi-C. This ambiguity is likely due to variability within the cellular population. Interrogating the mispredictions more, we found that, indeed, the mispredicted regions in GM12878 are highly variable in compartment status between independent replicates. Altogether these observations suggest that CoRNN is highly accurate for most bins, but that some sites are difficult to predict thanks to an ambiguous compartment status either due to unexpected chromatin marks or variability in the sampled population.

In our perturbation analysis, we also show that accurate predictions can be made using as few as two histone modification ChIP-seq datasets. This opens the possibility of predicting A/B compartments from cell lines and tissues where Hi-C data is not available. Since the ChIP-seq data is easier to obtain, cheaper, and more abundant, we envision that one can provide A/B compartment labeling for the entire ENCODE catalog of cell lines and tissues, especially with the help of imputed ChIP-seq signals (Schreiber et al., 2020a).

## ACKNOWLEDGMENTS

We would like to thank ENCODE Consortium’s Nuclear Architecture Working Group for helpful discussions and suggestions. This work was funded by National Institutes of Health awards U24 HG009446 and R35 HG011939.

## SUPPLEMENTARY MATERIAL

### Details for correct compartment value assignments

We calculated the correlation coefficient between the compartment values and the H3K4me3 ChIP-seq signals to correct the signs of the compartments. H3K4me3 has been observed to be positively correlated with the A/B compartment values (Lieberman-Aiden et al., 2009). Therefore, we flipped the sign of the compartments if the correlation coefficient was negative.

### Supplementary Tables

**Table 1.**
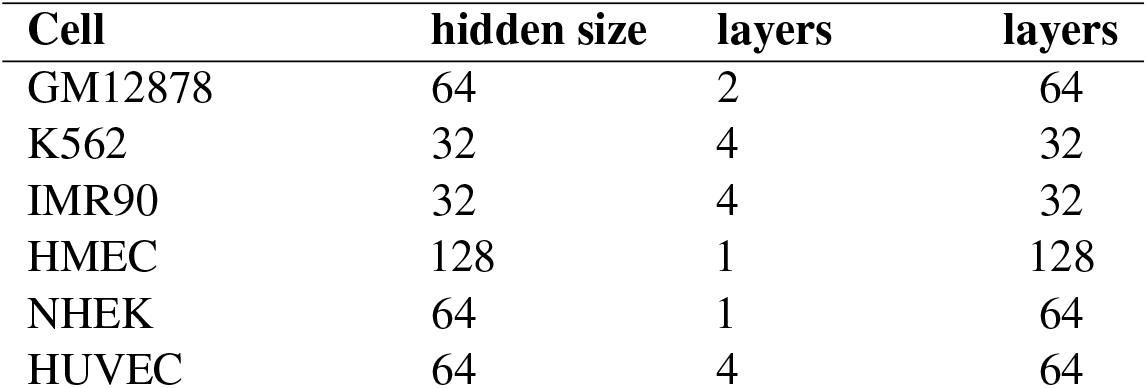
Hyper-parameters of CoRNN. For all cells, we trained the model using a batch size of 64, learning rate of 0.001, and 20 epochs.

| Cell | hidden size | layers | layers |
| --- | --- | --- | --- |
| GM12878 | 64 | 2 | 64 |
| K562 | 32 | 4 | 32 |
| IMR90 | 32 | 4 | 32 |
| HMEC | 128 | 1 | 128 |
| NHEK | 64 | 1 | 64 |
| HUVEC | 64 | 4 | 64 |

**Table 2.** Data summary of the six selected cells.

| Cell | A count | B count |
| --- | --- | --- |
| HUVEC | 13265 | 13714 |
| HMEC | 13140 | 13779 |
| IMR90 | 12691 | 14261 |
| GM12878 | 12754 | 14210 |
| K562 | 13099 | 13851 |
| NHEK | 14233 | 12707 |
| Total | 79182 | 82522 |

### Supplementary Figures

**Figure 8.**
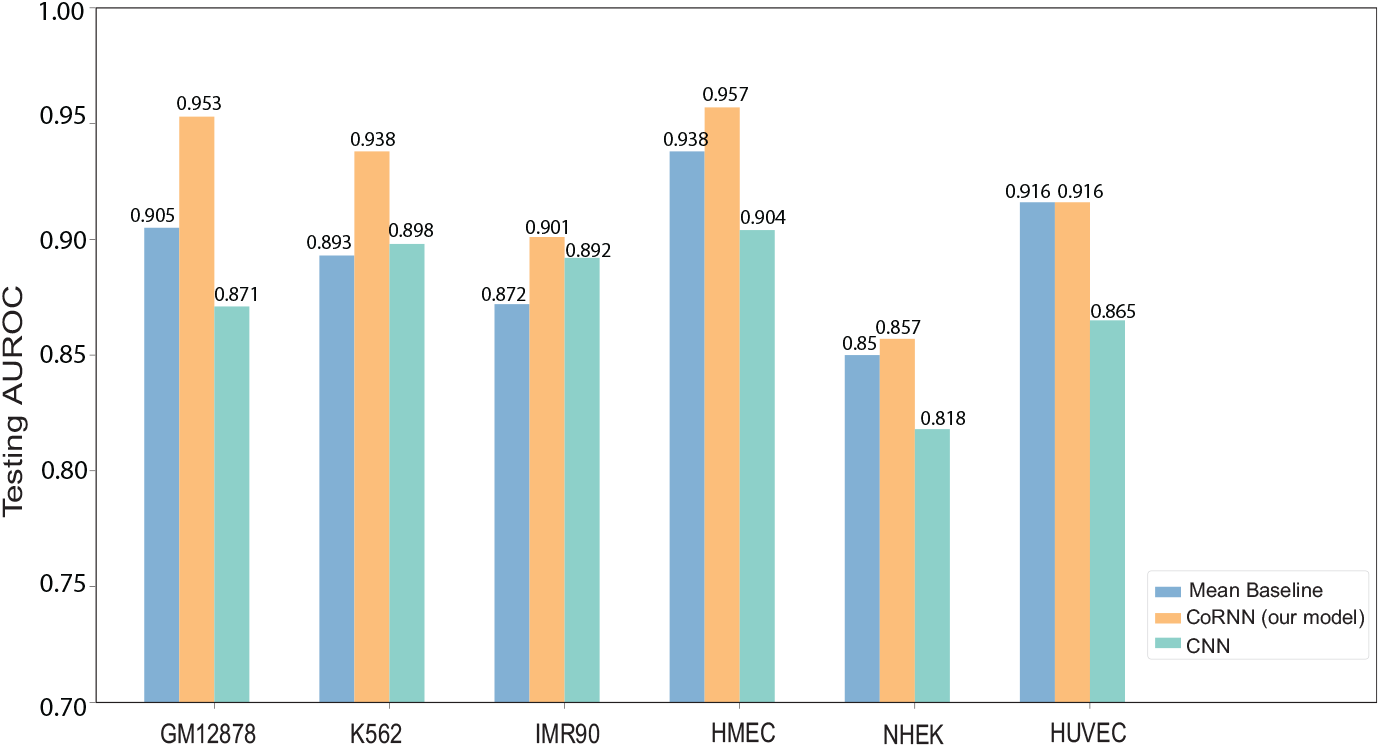
Testing results of CoRNN and Convolutional neural network model. Our model outperforms the convolutional neural network, thus justifying the choice of GRU as our neural network architecture.

**Figure 9.**
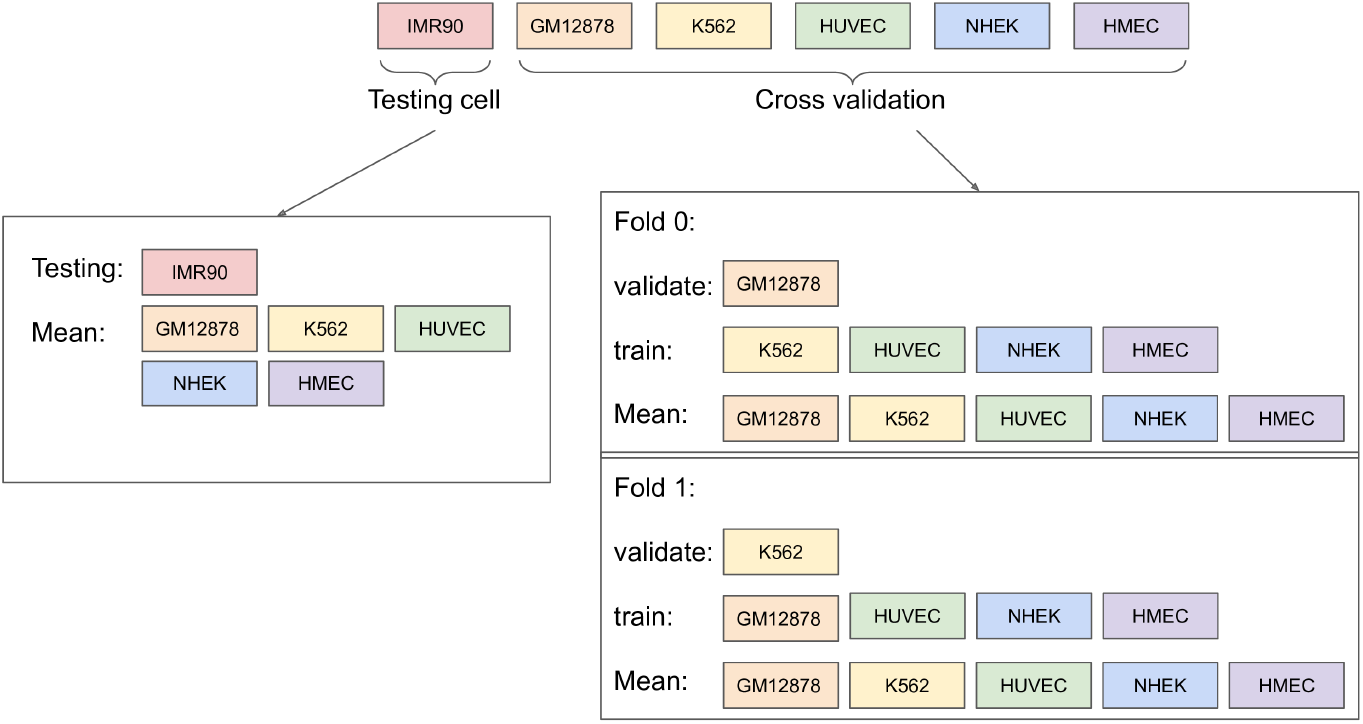
Cross-validation scheme for training CoRNN with IMR90 as the test cell line. Only creation of validation folds 0 and 1 are shown as examples. Similar process was used for creating both test and validation folds with other cell lines.

**Figure 10.**
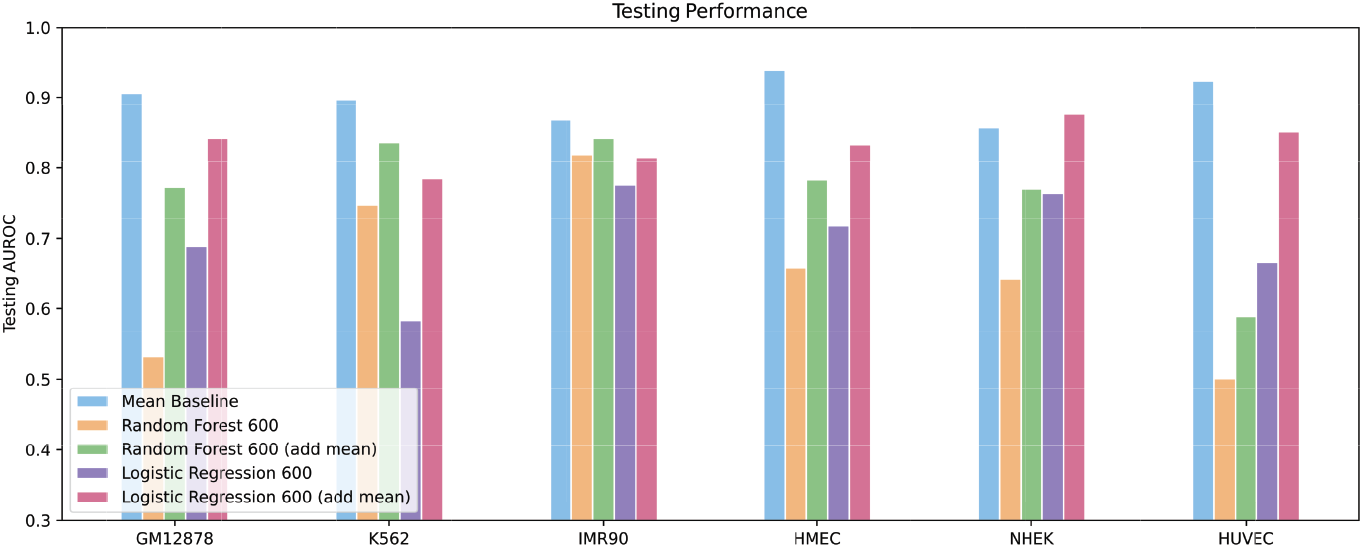
Random forest and linear regression models with concatenated 600 histone modification signal values as input. These models gave much worse performance than mean baseline. Therefore, we used these models with mean values of histone modification signals as inputs for baseline comparison.

**Figure 11.**
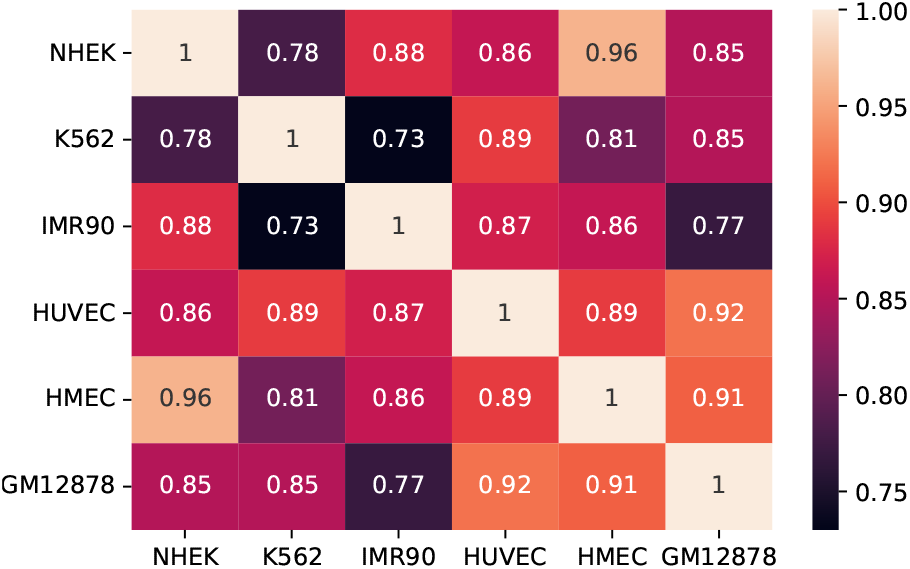
Correlation of compartment values across all six cell lines. Compartment values of HMEC, NHEK, and HUVEC have high correlations compared to the GM12878, K562, and IMR90 cell lines. This observation suggests that these cell lines are easier to predict using the *mean compartment value* baseline.

**Figure 12.**
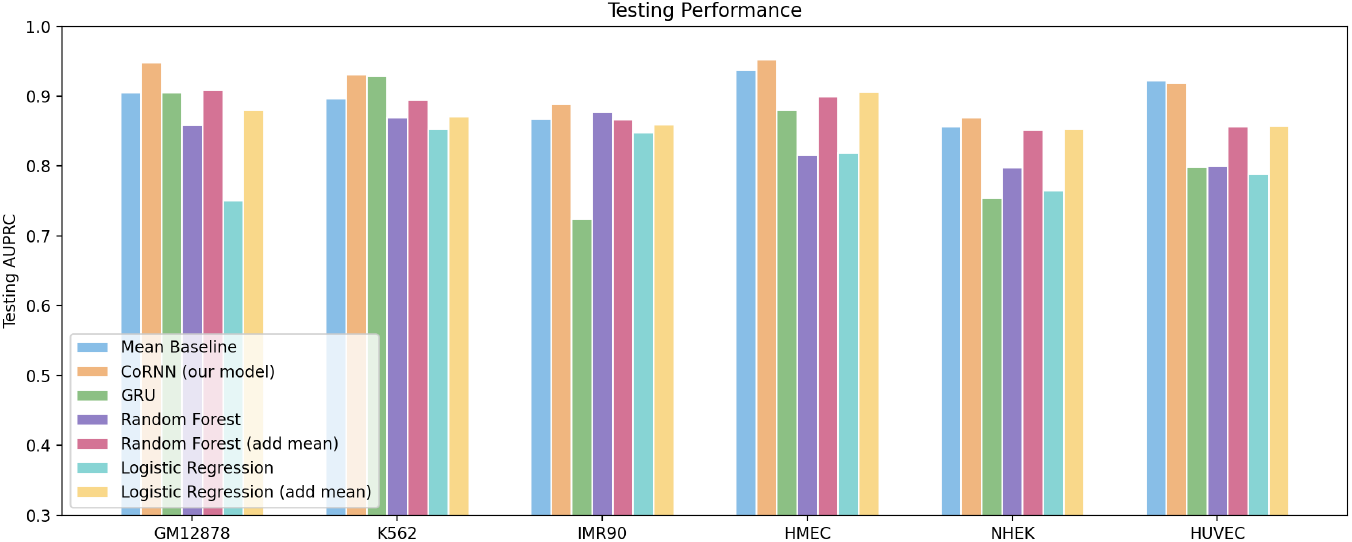
Testing results of CoRNN and baselines using AUPR score. Our model outperforms the mean baseline for five out of six cell lines

**Figure 13.**
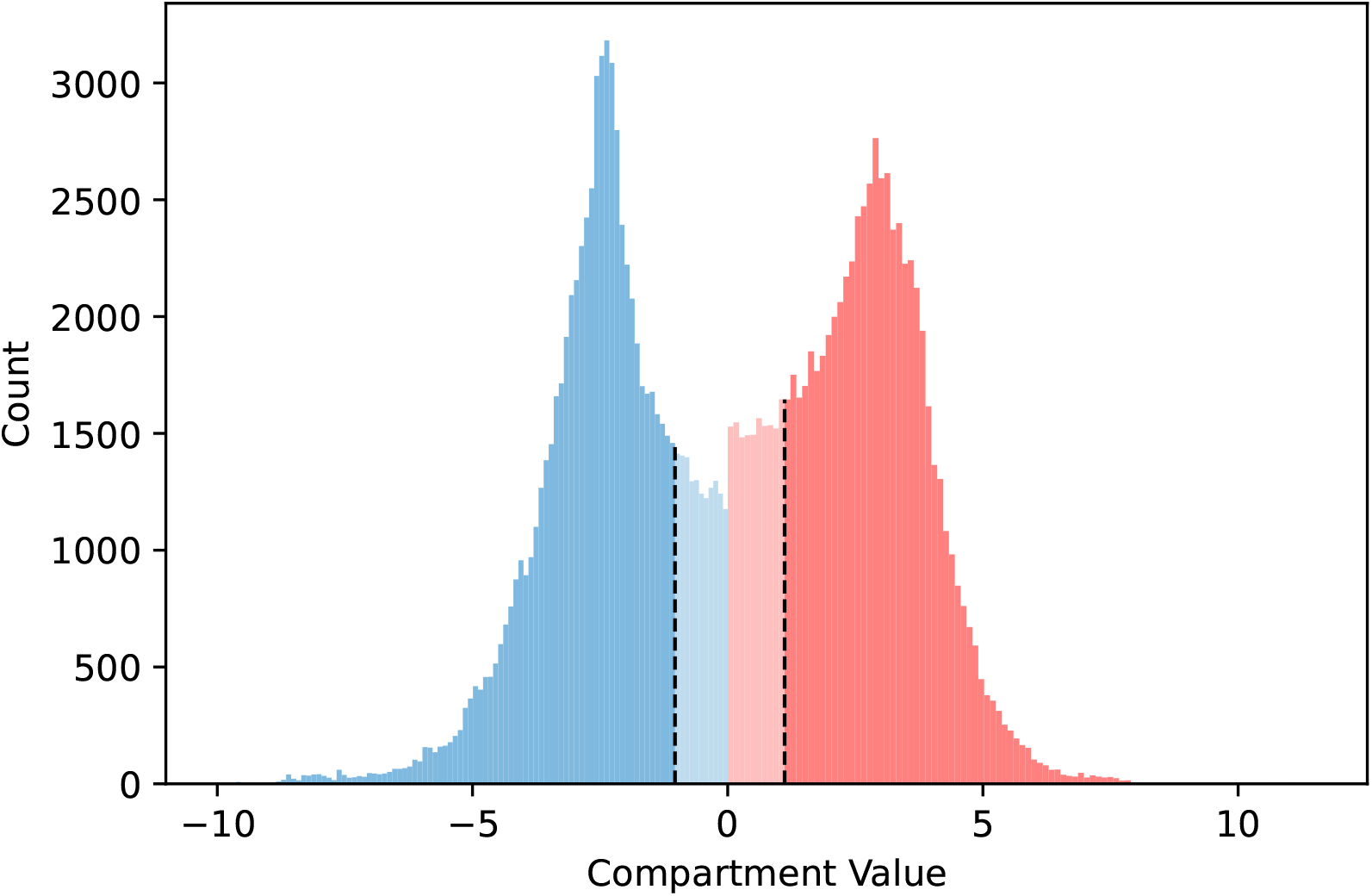
Picking strong compartments. We select strong compartments as those with absolute values > (*mean – std.deviation*) for all the compartment values.

**Figure 14.**
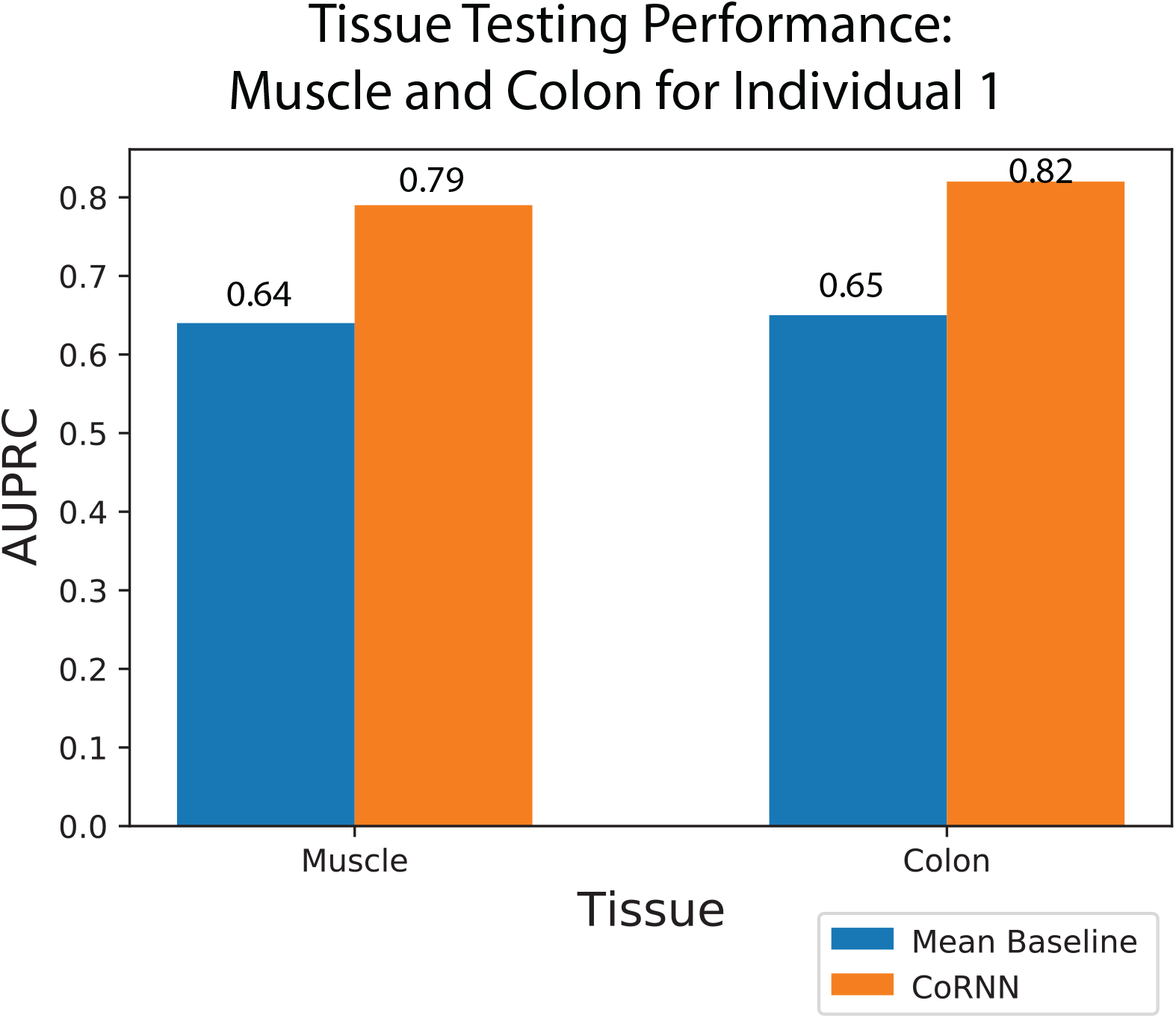
Testing performance of CoRNN and mean baseline on independent human muscle and colon tissue samples measured in AUPRC. CoRNN predicts A/B compartments for both tissues with higher cores compare to mean baseline.

